# Role of the NO-GC/cGMP signaling pathway in platelet biomechanics

**DOI:** 10.1101/2023.04.28.538670

**Authors:** Johanna G. Rodríguez, Aylin Balmes, Jan Seifert, Daniel Pinto-Quintero, Akif A. Khawaja, Marta Boffito, Maike Frye, Andreas Friebe, Michael Emerson, Francesca Seta, Robert Feil, Susanne Feil, Tilman E. Schäffer

## Abstract

Cyclic guanosine monophosphate (cGMP) is a second messenger produced by the NO-sensitive guanylyl cyclase (NO-GC) enzyme. In platelets, the NO-GC/cGMP pathway inhibits aggregation. One aspect of the inhibitory mechanism involves changes in the cytoskeleton; however, the molecular mechanisms underlying platelet inhibition and its correlation with cytoskeletal cellular stiffness are poorly understood.

We measured the cellular stiffness of individual platelets after treatment with the NO-GC stimulator riociguat or the NO-GC activator cinaciguat, using scanning ion conductance microscopy (SICM). We quantified changes in platelet shape using deep learning-based platelet morphometry. Cytoskeletal actin polymerization and platelet activation were measured by co-immunostaining F-actin and P-selectin, respectively. To test for clinical applicability of NO-GC stimulators in the context of increased thrombogenicity risk, we investigated the effect of riociguat on platelets from human immunodeficiency virus (HIV)-positive patients taking abacavir sulphate (ABC)-containing regimens, compared with HIV-negative volunteers.

Stimulation of human and murine platelets with the NO-GC stimulator riociguat or with the NO-GC activator cinaciguat downregulated P-selectin expression, decreased F-actin polymerization, and decreased cellular stiffness by ≈50%, compared to vehicle control. In addition, platelets became more circular, indicating decreased activation. Riociguat did not cause any change in platelet aggregation or circularity in HIV-positive patients taking ABC-containing regimens.

These results corroborate a functional role of the NO-GC enzyme in platelet biomechanics (cellular stiffness) in correlation with the inhibition of platelet activation and morphological changes. The observed changes in stiffness and platelet shape therefore demonstrate the possibility of pharmacologically targeting the NO-GC/cGMP pathway.

## 1. Introduction

Platelets are small anucleate blood cells responsible with critical roles in hemostasis and thrombosis [1]. Novel approaches to target inhibition of platelet activation, and hence thrombus formation, are currently being studied to reduce mortality in individuals affected by atherosclerosis, increased arterial stiffness [2], thromboembolism [3], or bleeding disorders such as thrombocytopenia [4]. Upon activation, platelets release inflammatory cytokines and become highly adhesive with a drastic change in shape [2]. The platelet cytoskeleton supports platelet shape in the resting state, and it changes dynamically during platelet activation in response to vascular damage [1]. Studies have shown that abnormalities in the biomechanical properties of platelets are related to inherited platelet cytoskeletal diseases [5]. Therefore, measuring platelet shape or cellular stiffness (amount of resistance of an object against deformation in response to an applied force), could help in the assessment of the bleeding risk in patients with cytoskeletal disorders [6].

Nitric oxide (NO) is an endogenous platelet inhibitor that binds to NO-sensitive guanylyl cyclase (NO-GC) [7]. In platelets, NO-GC has been shown to be the only NO receptor [8]. NO-GC catalyzes the conversion of guanosine 5’-triphosphate (GTP) to cyclic guanosine 3’,5’-monophosphate (cGMP) [9]. An increase in cGMP activates the cGMP-dependent protein kinase (PKG), which phosphorylates, among other proteins, vasodilator-stimulated phosphoprotein (VASP) in platelets and other cell types [10]. NO is involved in signaling cascades that support a healthy cardiovascular system including inhibition of platelet aggregation *in vitro* [9] and *in vivo* [11], making NO-GC an attractive target for cardiovascular pathologies including thrombosis [12]; however, continuous nitrate administration can lead to drug resistance because of the desensitization of NO-GC [7].

In a cardiovascular pathology such as ischemia, heme, the essential NO-GC co-factor, becomes oxidized, preventing NO from binding NO-GC and impairing cGMP generation [13]. Riociguat is a NO-GC stimulator, which has been approved for the treatment of pulmonary arterial hypertension and chronic thromboembolic pulmonary hypertension [14]. Riociguat has a dual effect on NO-GC: it stimulates the enzyme itself and decreases the dissociation of NO from NO-GC [15]. On the other hand, cinaciguat is a NO-GC activator that can activate NO-GC independently of its heme redox status (heme-oxidized or heme-free NO-GC) and irrespective of impaired NO signaling [16].

In the present study, we examined the effects of riociguat and cinaciguat on platelet biomechanics using scanning ion conductance microscopy (SICM), a non-invasive imaging technique that allows simultaneous measurement of morphological and mechanical properties in living cells [17,18] with high resolution. SICM has already been used to investigate the Young’s modulus (stiffness) of living migrating and non-migrating platelets [19] and to show that reduced platelet stiffness is related to increased bleeding in *MYH9*-related disease [20]. We correlated SICM measurements with F-actin and P-selectin (CD62P) immunostaining in human and wild-type (C57Bl6/J) or platelet-specific NO-GC knockout (KO) murine platelets. Additionally, platelet morphology was investigated with deep learning platelet morphometry [21].

Lastly, we studied the effect of riociguat on platelet morphology (circularity) in HIV-positive patients taking the anti-viral drug abacavir-sulphate (ABC) as part of their daily antiretroviral regimen, compared with platelets from HIV-negative volunteers. Currently, the first line of antiretroviral therapy for HIV-positive patients involves a combination of nucleotide reverse transcriptase inhibitors such as ABC [22]. Carbovir tri-phosphate (CBV-TP), the active ABC anabolite [22] is a guanosine derivative shown to compete and antagonize cGMP in the NO-GC/cGMP signaling pathway in platelets [22]. Therefore, the potential blockage of NO-GC/cGMP pathway by CBV-TP could increase the risk of thrombogenic events and lead to an increased development of cardiovascular pathologies in HIV patients taking the drug [22]. Our study is the first to analyze the effect on morphology (circularity) of the NO-GC stimulator riociguat in the setting of HIV.

## 2. Materials and Methods

### Animals

Platelets from 3-12 months old wild-type (C57Bl6/J) and megakaryocyte/platelet specific NO-GC KO mice with a Pf4-Cre (wt/Cre); NO-GC (flox/flox) genotype were used [23]. NO-GC KO mice were generated by crossing the Pf4-Cre line (B6-Tg(Cxcl4-cre)Q3Rsko/J) [24] to NO-GC(flox/flox) line (B6.129-Gucy1b3tm1.2Frb) [25]. Experiments were approved by the local authority (Regierungspräsidium Tubingen, IB 02/20M), and are reported in accordance with the ARRIVE guidelines [26].

### Isolation of human platelets

All procedures were approved by the institutional ethics committees (Medical Faculty and University Clinics at the University of Tübingen, 273/2018BO2 and 064/2022BO2 and Imperial College London Research Ethics Committee) in accordance with the declaration of Helsinki. Informed consent was obtained from all participants. Venous blood was collected from the antecubital vein of healthy volunteers in acid-citrate-dextrose (ACD) anticoagulant at 1:4 ratio (ACD: blood) at RT. Monovettes containing 4.8 mL of blood were centrifuged at 200 *× g* for 20 min at RT. Tyrode-HEPES (6 mM HEPES, 136.8 mM NaCl, 8.4 mM NaH_2_PO_4_, 2.6 mM KCl, 5.5 mM D-glucose, pH adjusted to 6.5 with HCl) was added to the platelet rich plasma (PRP) at a ratio of 1:3 and centrifuged at 920 *× g* for 10 min, without brake, at RT. The supernatant was discarded, and the platelet pellet was carefully re-suspended in 1 mL of Tyrode-HEPES, pH 7.4 and immediately used for experiments described below.

In addition, platelets from HIV-positive patients taking ABC-containing regimens registered at Chelsea and Westminster Hospital NHS Trust were obtained in accordance with research ethics permits 294707 21/NW/0148 approved by the NHS Health Research Authority and Chelsea and Westminster Hospital NHS Trust. All experiments were performed in accordance with the Declaration of Helsinki. Briefly, blood was collected from consented HIV-negative volunteers and HIV-positive patients taking ABC-containing regimens by venipuncture in VACUETTE tubes (Greiner Bio-One Ltd, United Kingdom) containing sodium citrate (3.2%). Whole blood was centrifuged at 175 *× g* for 15 minutes to obtain PRP. Washed platelets were obtained by addition of 150 μL of citrate-dextrose solution (C3821, Sigma-Aldrich, Dorset, UK) and 5 μL of prostaglandin E1 (P5515; Sigma-Aldrich, Dorset, UK) to the PRP, mixing by inversion and centrifuging at 1400 *× g* for 10 min. Platelet poor plasma was discarded, and the platelet pellet was re-suspended in a total volume of 20 mL of Tyrode’s-HEPES buffer (10 mM HEPES, 140 mM NaCl, 5 mM KCl, 1 mM MgCl_2_, 5 mM glucose, 0.42 mM NaH_2_PO_4_, 12 mM NaHCO_3_, pH 7.4). Then 3 mL of ACD and 5 μL of PGE_1_ were added and the tube was centrifuged again at 1400 *× g* for 10 min. Washed platelets were obtained by re-suspending the pellet in 1 mL of Tyrode’s-HEPES buffer and immediately used for experiments.

### Isolation of murine platelets

Murine blood was withdrawn from the retro-orbital sinus of wild-type and megakaryocyte/platelet-specific NO-GC KO mice under isofluorane anesthesia into a capillary with ACD buffer (85 mM sodium citrate, 72.9 mM citric acid, 110 mM D-glucose). 300 μl of platelet wash buffer (PWB) solution (4.3 mM K_2_HPO_4_, 4.3 mM Na_2_HPO_4_, 24.3 mM NaH_2_PO_4_, 5.5 mM D-glucose, 113 mM NaCl, pH 6.5), supplemented with 0.1% bovine serum albumin (BSA), were added into a plastic test tube where the blood was collected and then centrifuged at 200 *× g* for 2 min, without brakes, at RT. Collected PRP was transferred into a fresh tube while 600 μL of PWB solution were added to the remaining blood and centrifuged for a second time, with the same settings. Platelet supernatants from the two centrifugations were combined, and the PRP obtained after centrifugation at 2000 *× g* for 1 min, without brakes, at RT. Murine platelets were re-suspended in 600 μL Tyrode-HEPES buffer (10 mM HEPES, 137 mM NaCl, 12 mM NaHCO_3_, 2.7 mM KCl, 5.5 mM D-glucose, pH 7.4; supplemented with 0.1% BSA) and immediately used for experiments described below.

### Drug treatments for human and murine platelets

Thirty-five mm cell culture dishes with a glass bottom (81218, ibidi, Gräfelfing, Germany) were coated with 0.1 mg/mL fibrinogen (F3979, Sigma Aldrich, St. Louis, MO, USA) for 30 min at 37 °C. Tyrode-HEPES buffer was supplemented with CaCl_2_ (1mM) (10043-52-4, Sigma Aldrich) and MgCl_2_ (1 mM) (7786-30-3, Sigma Aldrich) to wash the platelets after spreading. NO-GC stimulator riociguat (10 μM) (9000554, Cayman, Ann Arbor, MI, USA), and NO-GC activator cinaciguat hydrochloride (10 μM) (SML1532, Sigma Aldrich), [1,2,4]oxadiazolo[4,3-a]quinoxalin-1-one (ODQ) (10 μM) (495320, Sigma-Aldrich, Dorset, UK), carbovir-5’-triphosphate triethylammonium salt (CBV-TP) (50 μM) (443952, US Biologicals, Salem, MA, USA) were dissolved in dimethyl sulfoxide (DMSO) (472301, Sigma Aldrich). 8-Br-cGMP (1 mM) (B1381, Sigma Aldrich) and adenosine 5’-diphosphate sodium salt (ADP) (3 μM) (20398-34-9, Sigma Aldrich) were dissolved in phosphate buffered saline (PBS) before adding to platelets for 10 min at 37 °C.

For SICM measurements, immunostaining quantification, and morphometric analysis, 50 μL of pre-treated platelets were added to the fibrinogen-coated cell culture dishes containing human or murine supplemented Tyrode-HEPES buffer and used as controls. Platelet adhesion and spreading were allowed to occur for 15 min at 37 °C, followed by three washes with human or murine supplemented Tyrode-HEPES buffer to remove non-adherent platelets before imaging.

### SICM imaging

Fibrinogen-coated cell culture dishes were installed in two custom-built SICM setups [27]. Live imaging of platelets was done within 1 h from collection because of their short life span. Borosilicate nanopipettes with an inner radius of 90 nm were fabricated using a CO_2_-laser-based micropipette puller (P-2000; Sutter Instrument, Novato, CA, USA). Topography and cellular stiffness images were obtained with a constant pressure of 10 kPa applied to the upper end of the pipette to allow for quantitative cellular stiffness measurements. The local cellular stiffness (Young’s modulus) was calculated from the slope of the ion current vs. distance curve (IZ-curve) between 99% and 98% of the saturation current (where the saturation current is the constant current at large tip-sample distances) for each pixel, using a model based on finite element calculation [28]. Imaging was done at a 25 Hz pixel rate, with 30 × 30 or 65 × 65 pixels at a scan size of 15 × 15 μm^2^. The cellular stiffness value of a single platelet was obtained by averaging the median value of each cellular stiffness map within the area of the platelet.

### F-actin and P-selectin (CD62P) immunostaining

Platelet F-actin and extracellular P-selectin (CD62P) expression, at baseline and after different drug treatments, were quantified by immunofluorescence. Briefly, platelets were fixed in PBS solution containing 2% formaldehyde (F1635, Sigma Aldrich) for 10 min and then washed three times with PBS. Fixed platelets were incubated with 1% BSA (A7906, Sigma Aldrich) in PBS for 10 min, and then with P-selectin (CD62P) primary monoclonal antibody (Psel.K02.PEeBioscience™, ThermoFisher) used at a 1:200 dilution, for 45 min at RT in a dark humidified chamber. Platelet F-Actin (1:2000 diluted in PBS) was stained with ActinGreen™ 488t (R37110, Invitrogen™, ThermoFisher) for 20 min at RT in a dark humidified chamber.

All epifluorescence images were recorded with an optical microscope (TiE, Nikon, Tokio, Japan) with a 100x/1.45 NA oil immersion objective and a monochrome digital camera (DS-Qi2, Nikon) with 300 ms exposure time. Images were then analyzed with ImageJ software (https://imagej.nih.gov/ij). Corrected total cell fluorescence (CTCF) for each individual platelet was obtained with the following formula: CTCF= fluorescence intensity integrated over the cell area – (mean fluorescence intensity of the background × cell area).

### Western blot analysis

Blood was collected from healthy humans, wild-type or genetically modified mice. The platelets were isolated and then re-suspended in human or murine Tyrode-HEPES buffer after isolation. DMSO (1:1000), 8-Br-cGMP (1 mM), riociguat (10 μM), cinaciguat (10 μM) were added to 100 μL of platelet suspension and pre-incubated for 10 min at 37 °C. Pre-treated platelets with the different drugs were centrifuged at 2000 *× g* for 1 min and then the supernatant was removed. Platelets were re-suspended in a lysis buffer (21 mM Tris pH 8,3; 0.67% SDS, 0.2 mM phenylmethylsulfonyl fluoride (PMSF) and 1 mL H_2_O), followed by freezing in liquid nitrogen and thawing. Protein lysates were separated by SDS-PAGE and proteins of interests detected with the following antibodies: phosphorylated (p-VASP) Ser239 (rabbit, 1:1000, Cell Signaling #3114S), glyceraldehyde-3-phosphate dehydrogenase (GAPDH) (rabbit, 1:1000, 14C10, Cell Signaling #2118S), NO-GC (sGC β1, rabbit, 1:100, abcam 24824).

### Microplate aggregation assay

PRP samples from HIV-negative volunteers and HIV-positive patients taking ABC-containing regimens were pre-incubated with riociguat (10 μM), ODQ (20 μM), DMSO (1:1000), or the different drug combinations for 30 min at 37 °C. PRP samples from HIV-negative volunteers were pre-incubated with CBV-TP (50 μM). An absorbance microplate reader (Sunrise, Tecan UK Ltd., UK) was used to read the light absorbance values in a 96-well plate (VWR; Leicestershire; UK) containing ADP (3 μM) and the pre-incubated PRP. Platelet aggregometry readings were performed at 20 s intervals for 16 min at 37 °C, shaking for 7 s before each reading. Maximum aggregation values at minute 16 were obtained by calculating the percentage change in comparison to the baseline.

### Deep learning platelet morphometry

Platelet circularity and area were analyzed using a convolutional neural network [21].Circularity is a unitless morphological parameter ranging between 0 and 1; a lower circularity indicates platelet shape irregularity, while a higher circularity represents a rounder shape [21]. Phase contrast images for platelet morphometry were recorded with an optical microscope (TiE, Nikon, Tokyo, Japan) on fixed platelets using a 100x/1.45 NA immersion oil objective for human and murine platelets. Phase contrast images of fixed and washed platelets from HIV-negative volunteers and HIV-positive patients taking ABC-containing regimens were taken at the FILM (Facility for Imaging by Light Microscopy) at Imperial College London. WF3 Zeiss Axio Observer, with a Zen Blue software with a 100x/1.4 Oil Ph3 Plan Apochromat and phase illumination for the transmitted light channel, and a Hamamatsu Flash4.0 camera with a pixel size of 65 nm were used to record the images.

### Statistics

Data were analyzed and processed in Igor Pro (WaveMetrics, Lake Oswego, Oregon, USA). Data are presented as median ± median absolute deviation (MAD) unless stated otherwise. All results were tested using Tukey’s test for parametric multiple comparisons, except for circularity data where Dunn’s test for non-parametric multiple comparisons was applied. Results are considered significantly different for *P-*values < 0.05.

## 3. Results

Platelet activation, stiffness, and shape changes were investigated in platelets isolated from human, wild-type, and platelet-specific NO-GC KO mice.

### Platelet activation is dampened by riociguat and cinaciguat

We first examined the effect of the NO-GC stimulator riociguat and the NO-GC activator cinaciguat on the cytoskeleton and activation for human platelets as well as platelets isolated from wild-type (C57Bl6/J) and platelet-specific NO-GC KO mice (Figure 1). We found that the cGMP-modulating drugs 8-Br-cGMP, riociguat, or cinaciguat decreased F-actin (Figure 1b, e) and P-selectin (Figure 1c, f) in isolated human platelets and wild-type murine platelets by ≈50%, compared to vehicle control (DMSO).

**Figure 1.**
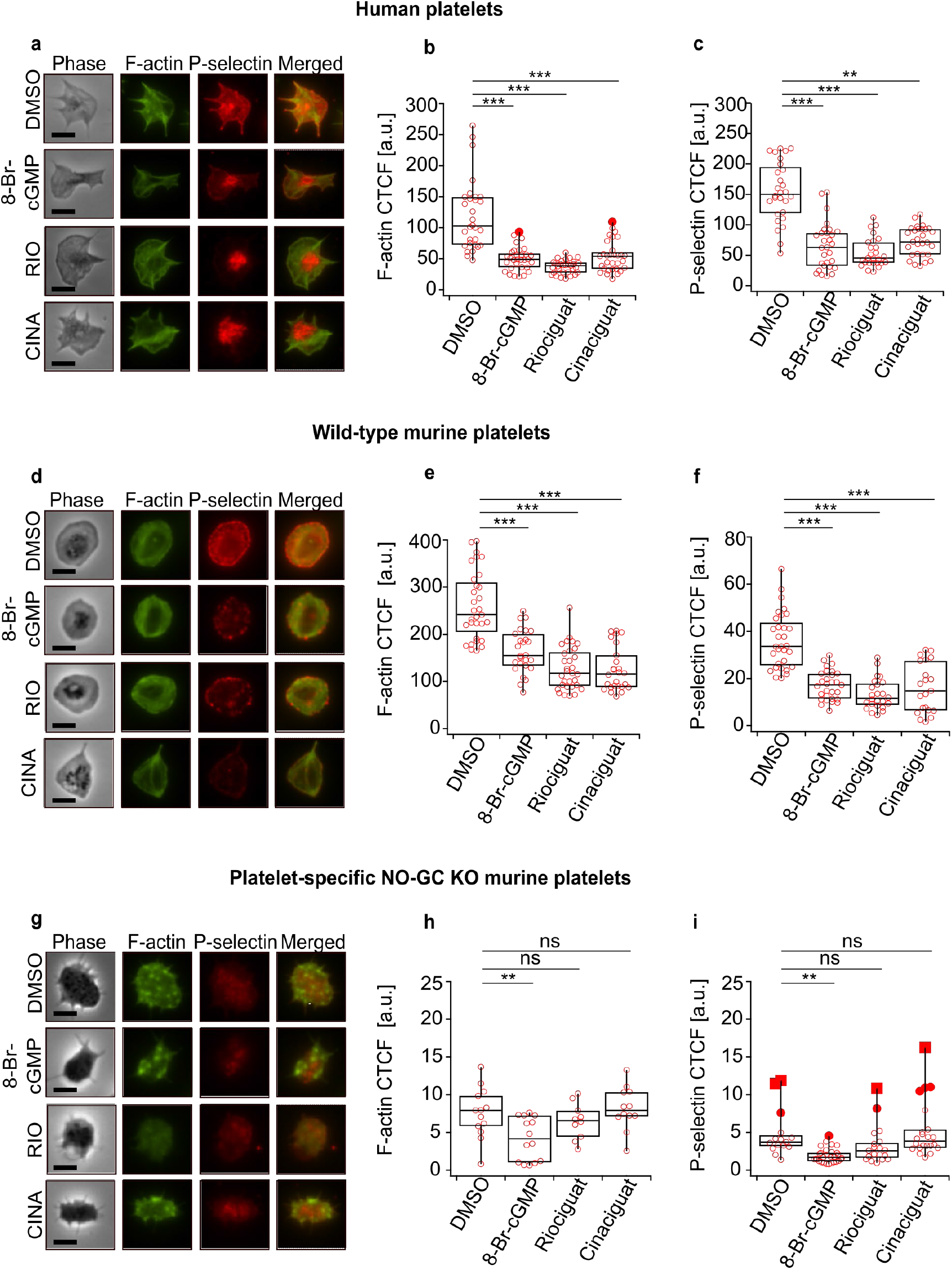
F-actin and P-selectin are decreased in human platelets and wild-type (C57Bl6/J) murine platelets when treated with 8-Br-cGMP, riociguat, or cinaciguat. **(a, d, g)** Co-staining of F-actin and P-selectin in washed human, wild-type murine, and platelet-specific NO-GC murine KO platelets. **(b, e, h)** F-actin and **(c, f, i)** P-selectin fluorescence quantification. DMSO (1:1000), 8-Br-cGMP (1 mM), riociguat (10 μM), cinaciguat (10 μM). The significance level of *P* values is indicated by asterisks (^*^ *P* < 0.05; ** *P* < 0.01; ^***^ *P* < 0.001; ns, not significant; Tukey’s test). Scale bars: 5 μm. Experiments were performed with ≈15 platelets per donor from *n*=2 (human), *n*=2 (wild type mice), *n*=2 (NO-GC KO mice) donors.

The P-selectin CTCF values ranged 2 – 225 [a.u.] in human platelets, 2 – 80 [a.u.] in wild-type murine platelets, and 2 – 25 [a.u.] in NO-GC KO murine platelets. Thus, these values indicate a lower basal protein expression of F-actin (Figure 1i) and P-selectin (Figure 1h) in the KO mice. Moreover, no further decreases in F-actin and P-selectin were observed in NO-GC KO platelets treated with riociguat or cinaciguat, whereas 8-Br-cGMP decreased F-actin and P-selectin by an additional ≈50% (Figure 1g, h, i). Downregulation of P-selectin suggests an inhibitory role of the NO-GC pathway in platelet activation (Figure 1c, f, i).

Interestingly, P-selectin was observed mostly in the form of nodule-like local spots that might be individual granules distributed throughout the platelet in the treated wild-type murine platelets (Figure 1d), while a more homogenous distribution was observed in the case of NO-GC KO murine platelets (Figure 1g). For human platelets, nodule-like spots were only observed after treatment with riociguat or cinaciguat (Figure 1a). Overall, these results indicate that F-actin and P-selectin are modulated by the NO-GC enzyme and that an increase in cGMP decreases the downregulation of F-actin and P-selectin. The NO-GC KO animals were used to show that the drug effect from riociguat or cinaciguat is indeed mediated by NO-GC.

### Platelet stiffness is decreased by riociguat and cinaciguat

SICM was used to investigate the role of NO-GC in the biomechanical properties (stiffness) of wild-type murine, platelet-specific NO-GC KO murine, and human platelets. Topography and stiffness images of platelets were recorded with high spatial resolution (Figure 2). Human (Figure 2b) and wild-type murine platelets (Figure 2e) treated with cGMP-modulating drugs had a significantly decreased average cellular stiffness (≈2 kilopascal, kPa) in comparison to DMSO-treated platelets (≈5 kPa), indicating a decrease of ≈50% in cellular stiffness.

**Figure 2.**
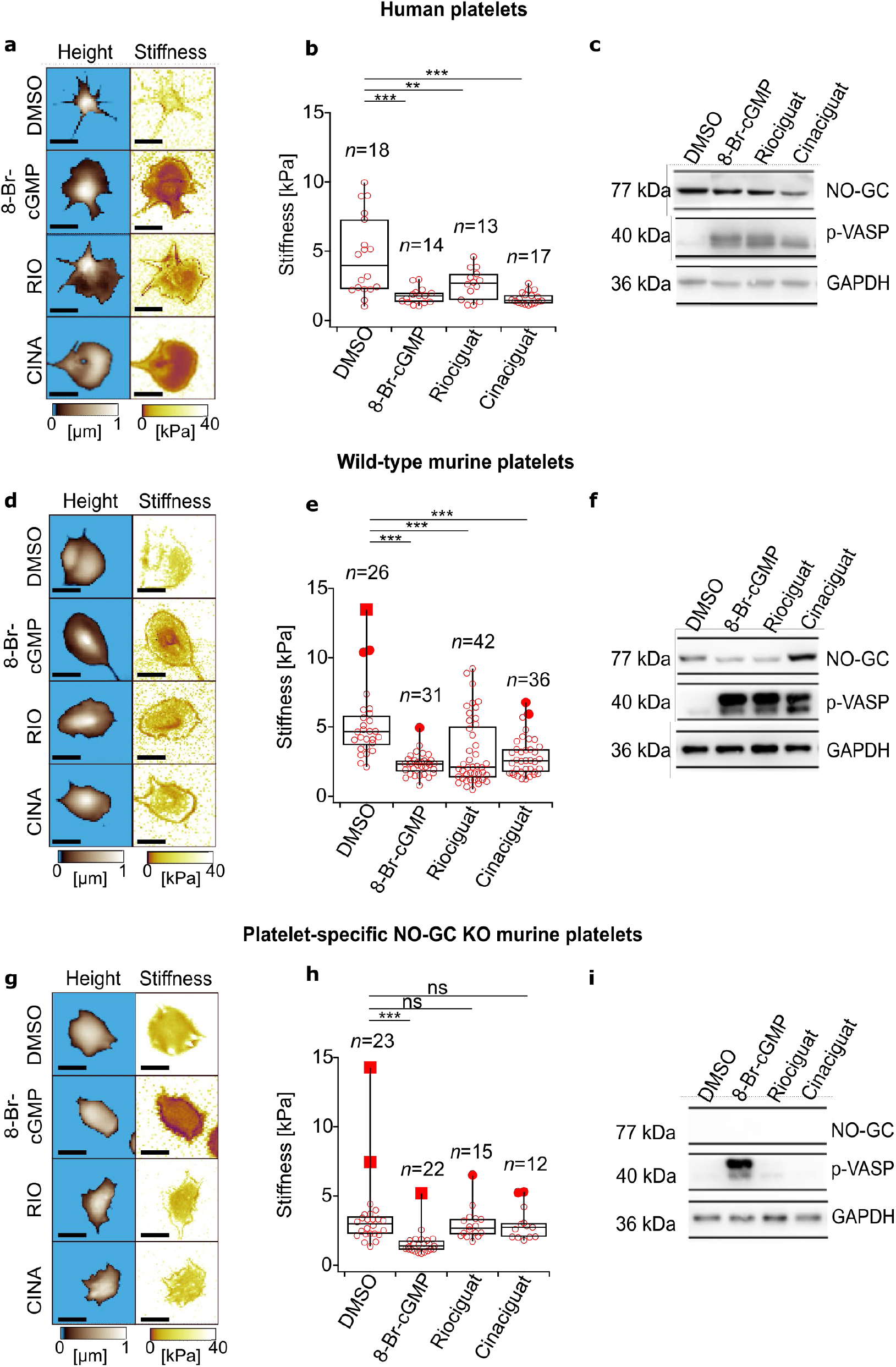
Cellular stiffness is decreased in human platelets and wild-type (C5BI6/J) murine platelets when treated with 8-Br-cGMP, riociguat or cinaciguat. **(a, d, g)**. Topography and cellular stiffness images of washed human platelets, wild-type murine, and platelet-specific NO-GC murine KO platelets. **(b, e, h)** Cellular stiffness quantification. **(c, f, i)** Western blot analysis of VASP phosphorylation (p-VASP) and NO-GC. GAPDH was used as loading control. DMSO (1:1000), 8-Br-cGMP (1 mM), riociguat (10 μM), cinaciguat (10 μM). The significance level of *P*-values is indicated by asterisks (^*^ *P* < 0.05; ^**^ *P* < 0.01; ^***^ *P* < 0.001; ns., not significant; Tukey’s test). Scale bars: 5 μm. Experiments were performed with 6-21 platelets per donor from *n*=2 (human), *n*=2 (wild type mice), *n*=2 (NO-GC KO mice) donors. Control blots were from the same sample, and performed from *n*=3 (human), *n*=3 (wild type mice), *n*=4 (NO-GC KO mice) donors.

In contrast, NO-GC KO platelets (Figure 2h) showed no decrease in cellular stiffness after the treatment with riociguat or cinaciguat (≈5 kPa) in comparison with DMSO (≈5 kPa). A change in cellular stiffness by ≈60% was only observed when NO-GC KO platelets were treated with 8Br-cGMP (≈2 kPa) (Figure 2b, c, d). Furthermore, we found increased levels of phosphorylated VASP (p-VASP), a downstream effector of cGMP and regulator of F-actin polymerization [29], in wild-type and human platelets for all the treatments (Figure 2c, f). In contrast, VASP phosphorylation was almost undetectable in NO-GC KO platelets treated with riociguat or cinaciguat, but still detectable when NO-GC KO platelets were stimulated with 8-Br-cGMP (Figure 2i).

Efficient NO-GC deletion in platelets of megakaryocyte/platelet-specific NO-GC KO mice was confirmed via Western blot (Figure 2i). Taken together, these results suggest that NO-GC-mediated VASP phosphorylation and actin cytoskeletal rearrangements (Figure 1) may be linked to the regulation of cellular stiffness in platelets (Figure 2). Specific deletion of NO-GC in platelets corroborated the role of NO-GC in platelet stiffness treated with riociguat or cinaciguat, suggesting that stiffness can be modulated through the NO-GC/cGMP pathway.

### Platelet shape is influenced by riociguat and cinaciguat

Platelet shape changes were investigated with deep learning platelet morphometry (Figure 3). A convolutional neural network (CNN) was used to generate binary prediction images from optical phase contrast images for the analysis of the circularity of human, murine wild-type, and platelet-specific NO-GC KO platelets. Platelet circularity could be an indicator of platelet activation and spreading stages. When human platelets and wild-type murine platelets were treated with cGMP-modulating drugs, a significant increase in platelet circularity was observed (Figure 3a-d). However, the circularity (circ) of platelets from platelet-specific NO-GC KO mice did not change when treated with riociguat (circ = 0.5; *P* = 0.98) or with cinaciguat (circ = 0.5; *P* = 0.94), compared to DMSO (circ = 0.5) (Figure 3e, f). The absence of a change in the circularity of NO-GC KO platelets treated with riociguat or cinaciguat demonstrated that platelet shape is likely modulated via NO-GC-mediated signaling.

**Figure 3.**
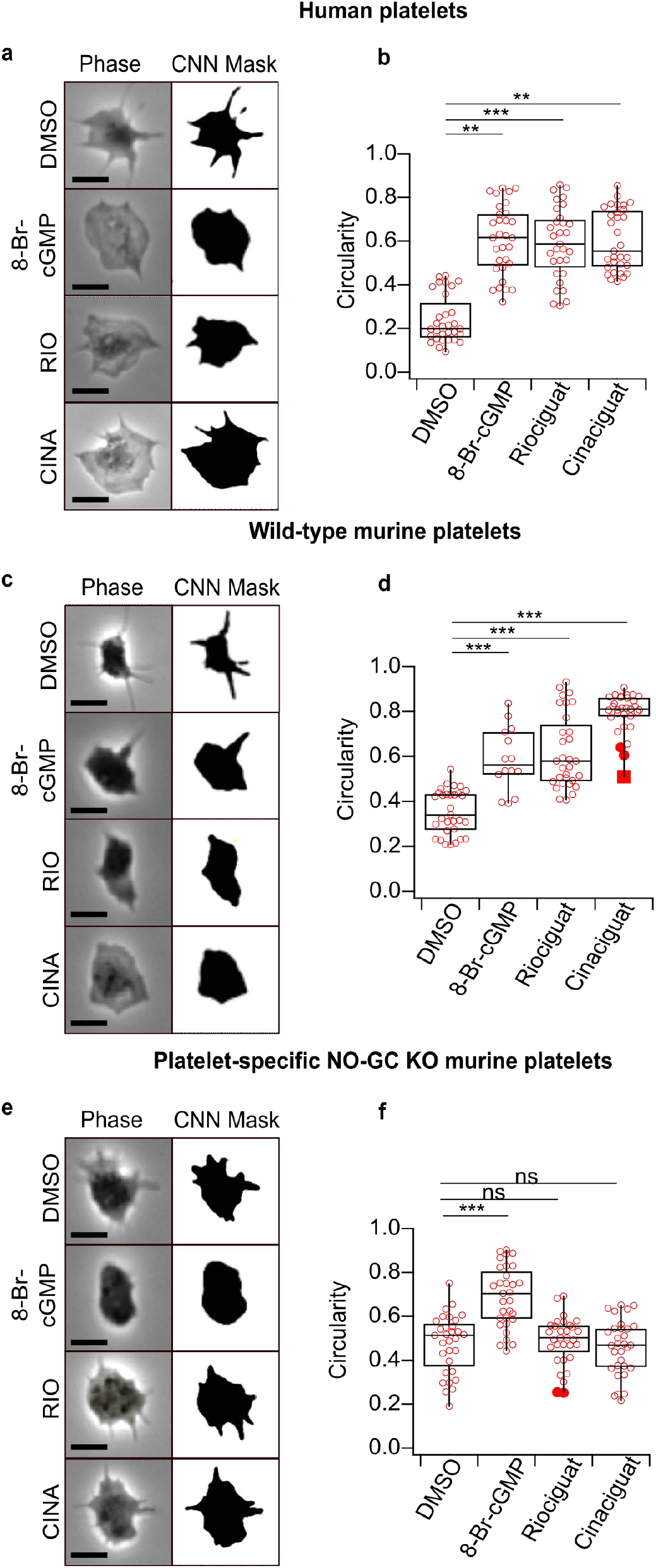
Platelet shape is altered in human platelets and wild-type (C57Bl6/J) murine platelets when treated with 8-Br-cGMP, riociguat, or cinaciguat. **(*a, c, e*)** Phase contrast and binary prediction images obtained via deep learning morphometry in washed human, wild-type murine, and platelet-specific NO-GC KO murine platelets. **(b, d, f)**. Platelet circularity. DMSO (1:1000), 8-Br-cGMP (1 mM), riociguat (10 μM), cinaciguat (10 μM). Platelet numbers for all conditions: *n*=30. The significance level of *P*-values is indicated by asterisks (^*^*P* < 0.05; ^**^*P* < 0.01; ^***^*P* < 0.001; ns, not significant; Dunn’s test). Scale bars: 5 μm. Experiments were performed with ≈15 platelets per donor from *n*=2 (human), *n*=2 (wild type mice), *n*=2 (NO-GC KO mice) donors.

### Circularity and aggregation are decreased in riociguat-treated platelets from HIV-negative donors but not in platelets from HIV-positive patients treated with ABC

The circularity of platelets from HIV-negative platelet donors was significantly increased (*P* = 0.001) by ≈50% when treated with riociguat (circ = 0.60) compared to DMSO (circ = 0.31) (Figure 4a), consistent with our findings of riociguat-treated platelets from healthy human volunteers (Figure 3b). However, the circularity of platelets from HIV-negative platelet donors did not change (*P* = 0.71) when treated with the active ABC anabolite CBV-TP (circ = 0.29), which mimics CBV-TP present in HIV-positive patients taking ABC-containing regimens, compared to DMSO control (circ = 0.31) (Figure 4a). Interestingly, the circularity of riociguat-treated platelets from HIV-positive patients taking ABC-containing regimens (circ = 0.37) did not change compared to DMSO control (circ = 0.34) (Figure 4b). Additionally, ADP median circularity for HIV-negative volunteers (circ = 0.21) and HIV-positive patients taking ABC-containing regimens (circ = 0.25) was ≈30 % smaller than for DMSO, albeit not significantly different (Figure 4a, b). These findings could indicate a disruption of the thrombo/cardio-protective NO-GC/cGMP signaling pathway in HIV-positive patients treated with ABC-containing regimens.

**Figure 4.**
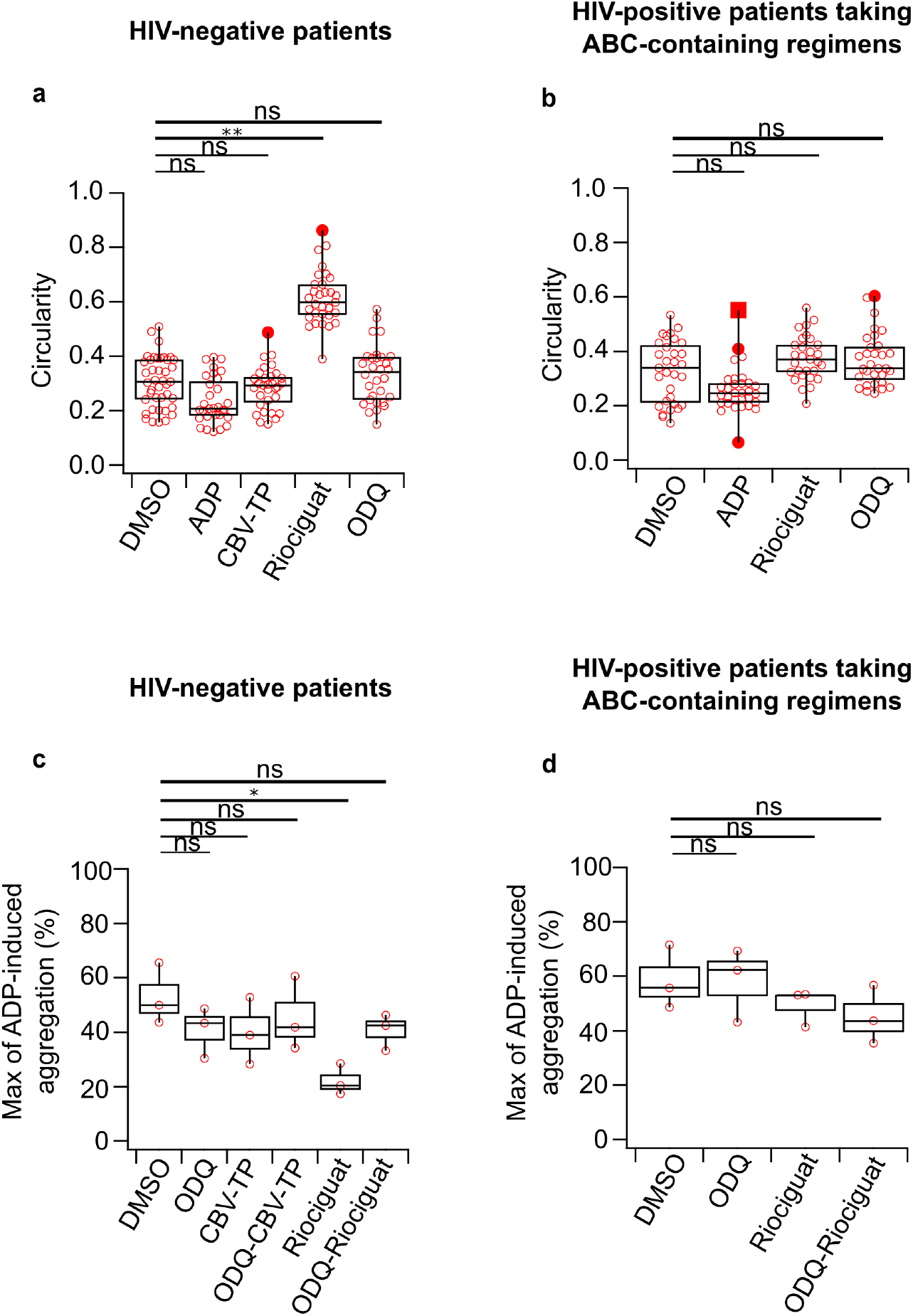
Riociguat increases platelet circularity and decreases platelet aggregation in HIV-negative volunteers, but not in HIV-positive patients taking ABC-containing regimens. **(a, b)** Platelet circularity. **(c, d)** Maximum of ADP-induced aggregation in response to indicated drug treatments. ADP (3 μM), DMSO (1:1000), CBV-TP (50 μM), riociguat (10 μM), ODQ (20 μM). Platelet numbers for circularity: *n* ≈ 15 per donor from *n*=2 donors. The significance level of *P-* values is indicated by asterisks (^*^ *P* < 0.05; ^**^ *P* < 0.01; ^***^ *P* < 0.001; ns, not significant; Tukey’s test). Donors for aggregation experiments: *n*=3.

Similarly, microplate reader-based analysis of platelet aggregation showed decreased aggregation of platelets isolated from HIV-negative donors when treated with riociguat (58% for riociguat, 20% for DMSO; *P* = 0.027) (Figure 4c), but not in platelets isolated from HIV-positive patients ABC-containing regimens (58% for both riociguat and DMSO; *P* = 0.58) (Figure 4d). Moreover, platelets isolated from HIV-negative volunteers treated with CBV-TP showed a similar percentage of aggregation as platelets from HIV-negative volunteers treated with the selective NO-GC antagonist ODQ (*P* = 0.99; percentage of aggregation= 42% to 40%) (Figure 4c). No pharmacological interaction of CBV-TP with ODQ was observed (Figure 4c). Likewise, ODQ showed no effect in platelet aggregation when pre-incubated together with riociguat in HIV-positive patients taking ABC-containing regimens (Figure 4d). Additionally, when comparing the different drug combinations (ODQ-CBV-TP for HIV-negative patients, ODQ-riociguat for both HIV-negative and HIV-positive patients taking ABC-containing regimens), the maximum ADP-induced aggregation was identical (≈42%), and no significant difference compared to aggregation values for DMSO (≈42%) was detected.

Taken together, our findings show that activation of the NO-GC/cGMP pathway in platelets results in platelet de-activation (or maintenance of platelets in an inactive state) and interestingly correlates with a decrease in cellular stiffness, downregulation of F-actin, and decrease in platelet circularity (Figure 5). Platelet shape changes (circularity increase) observed in riociguat-treated platelets from healthy volunteers were not observed in riociguat-treated platelets from HIV-positive patients currently taking ABC (Figure 4b). Riociguat did not exert any changes in platelet shape in platelets isolated from HIV-positive patients currently taking ABC, because the thrombo-protective effect of the NO-GC/cGMP pathway was abolished. Our data further suggest that CBV-TP could block the NO-GC/cGMP pathway and thereby the inhibition of platelet aggregation (Figure 5).

**Figure 5.**
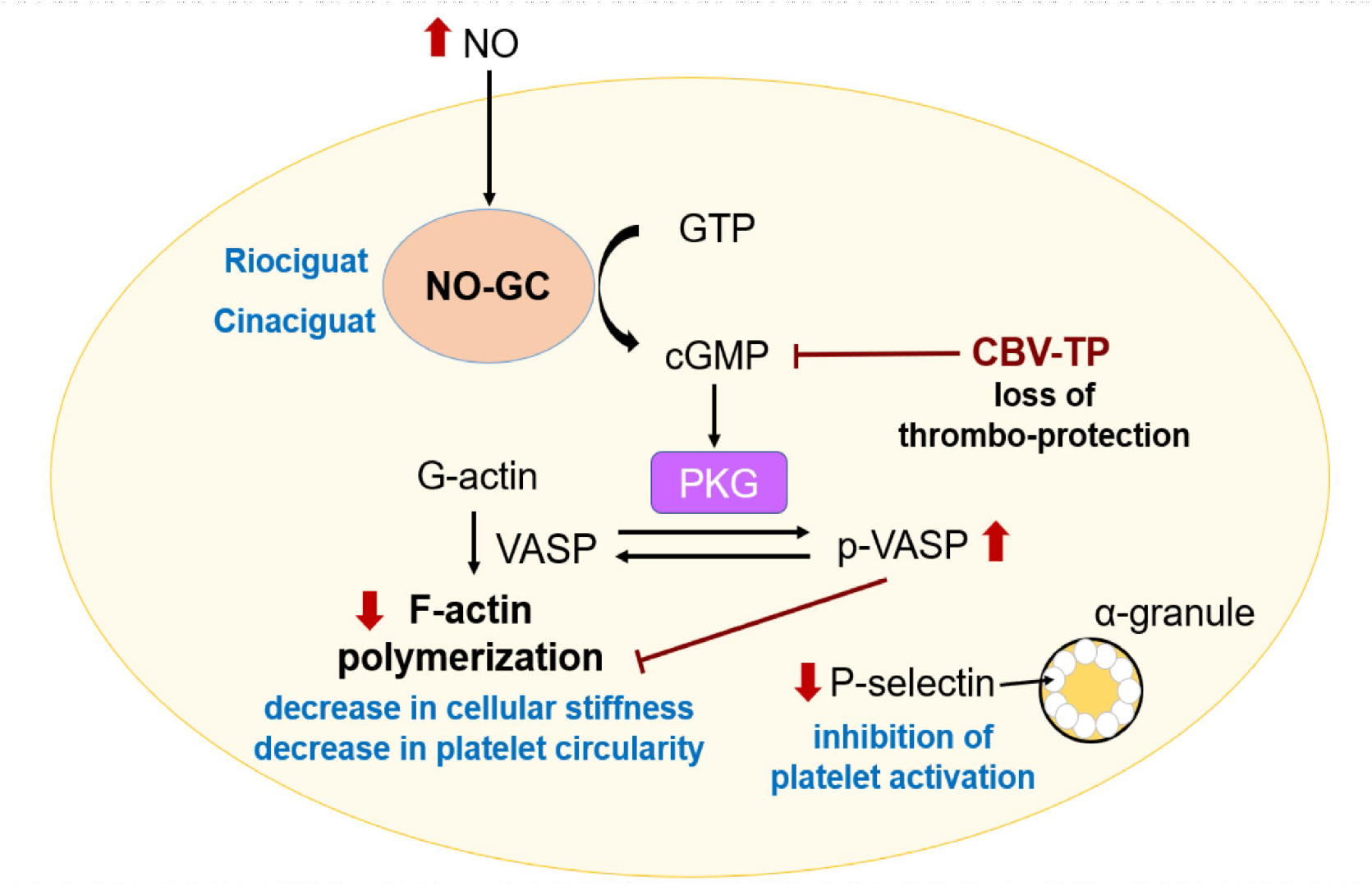
Role of the NO-GC/cGMP pathway in the inhibition of platelet activation, decrease in cellular stiffness, and loss of thrombo-protection.

## 4. Discussion

The goal of this study was to investigate molecular mechanisms that mediate platelet biomechanics, which could become novel strategies for the treatment or diagnosis of thrombosis and related cardiovascular pathologies in a variety of clinical settings. Specifically, we sought to elucidate the role of the NO-GC/cGMP signaling pathway in human and murine platelets, with a focus on cytoskeleton biomechanics.

Platelets store P-selectin in their alpha granules and upon platelet activation, the external membrane of the platelet exposes P-selectin [30]. We found that P-selectin expression and cytoskeletal F-actin polymerization were decreased in human and wild-type murine platelets after treatment with different cGMP-modulating drugs (Figure 1a and 1d) and these changes correlated with decreased platelet cellular stiffness (Figure 2a, 2d). The specificity of our findings was confirmed with platelets from platelet-specific NO-GC KO mice, where cGMP-modulating drugs did not have any effect on cytoskeletal F-actin (Figure 1h), P-selectin (Figure 1i), platelet cellular stiffness (Figure 2g), or platelet circularity (Figure 3f).

At the molecular level, we also observed an increase in phosphorylated VASP (Figure 2c, 2f). Phosphorylation of VASP at serine239 (p-VASP) is a marker for PKG activity and NO-mediated effects in platelets [31] and is involved in the downregulation of F-actin polymerization [32], which leads to a decrease in platelet cellular stiffness. The phosphorylation of VASP in platelets is also associated with inhibition of fibrinogen receptor (integrin GPIIb/IIIa) [33]. Fibrinogen binding to GPIIb/IIIa has been linked to thrombus stabilization [34], suggesting that the activation of NO-GC/cGMP signaling and subsequent VASP phosphorylation decreases thrombus formation [10].

Platelet shape (circularity) is mediated by actin-binding proteins [32], and upon platelet activation, filopodia and lamellipodia are formed. Platelet shape change is essential for platelet adhesion [35] and spreading, which occur via four different stages [36]: (1) platelet adhesion; (2) formation of filopodia; (3) development of filopodia and lamellipodia; and (4) presence of lamellipodia and of orthogonally arrayed short actin filaments inside the lamellipodia [36]. Our results suggest that the circularity in platelets treated with 8-Br-cGMP, riociguat, or cinaciguat is significantly increased (Figure 3b, 3d), owing to a more circular platelet spreading state and shape with fewer filopodia and lamellipodia. This gives an indication that NO-GC plays a role in platelet shape change, because the circularity was not altered by the treatments in NO-GC KO platelets (Figure 3f). Platelet morphological features such as circularity could define morphological subtypes such as not-spread, partially, or fully spread [37]. According to the circularity values obtained in this study, most of the platelets that were stimulated with cGMP stimulating drugs fit within a partially spread category. This suggests that when platelets increase cGMP upregulation through NO-GC stimulation (riociguat) or activation (cinaciguat), platelets do not reach a fully spread state or become fully activated. NO-GC has a basal, NO-independent activity, which would add to the NO-induced, endothelium-derived inhibition and probably influence platelet morphology (roundish shape).

Similarly, for platelets treated with riociguat, increased circularity values were observed in healthy volunteers but not in HIV-positive patients taking ABC (Figure 4), suggesting that the NO-GC/cGMP pathway may be downregulated or blocked by off-target effects of antiretrovirals such as ABC in these patients. This thinking is supported by the fact that platelet circularity was not altered by any GC-modulating treatment suggesting platelets are hyperactivated in these patients. Consistent with this finding, riociguat decreased platelet aggregation in HIV-negative volunteers but not in HIV-positive patients currently taking ABC regimens (Figure 4e, 4d), suggesting that CBV-TP, the active ABC anabolite, inhibits the NO-GC/cGMP pathway in platelets. However, further studies on the interaction between cGMP and ABC are warranted. Our studies suggest that treatment of HIV-positive patients with ABC-containing regimens blocks NO-GC/cGMP-mediated thrombo-protection (readout circularity), potentially putting these patients at higher risk of thrombotic events. This is supported by observational studies, indicating an increased risk of myocardial infarction in patients taking ABC [38].

Platelets have high concentrations of PKG (above 0.1 μmol/mL), mainly the PKG-Iβ isoform [39]. Platelets can also be regulated by phosphodiesterases (PDEs) [40] through the hydrolysis of cyclic adenosine 3’, 5’-monophosphate (cAMP) and cGMP [41]. Dipyridamole is an unspecific PDE inhibitor that decreases cGMP hydrolysis, and it is occasionally used in combination with acetylsalicylic acid in therapy to inhibit platelet activation; however, its efficacy is questioned [39]. Platelet inhibition occurs via cGMP/PKG activation, which is consistent with our finding that in platelets lacking NO-GC, treatments with NO-GC stimulator or activator did not have any effect on the platelet activation marker P-selectin (CD62P), suggesting that cGMP/PKG is essential for platelet inhibition.

In summary, our study indicates that the NO-GC/cGMP signaling pathway is involved in the inhibition of P-selectin expression and downregulation of cytoskeletal F-actin polymerization. Moreover, when cGMP/PKG is stimulated via a NO-GC stimulator (riociguat) or a NO-GC activator (cinaciguat), a decrease in platelet cellular stiffness and platelet circularity occurs via NO-GC. Furthermore, we conclude that NO-GC/cGMP signaling regulates the cellular stiffness and shape changes of platelets. As cellular stiffness indicates the mechanical state of the platelet cytoskeleton, measuring cellular stiffness could become a novel biomarker to evaluate patients’ cardiovascular risk, considering that platelet hyperactivation is associated with multiple cardiovascular conditions such as increased arterial stiffness [42], detrimental blood flow [43], or atherosclerosis. Also, platelet circularity could give a hint of platelet activation state suggesting that circularity could also become a biomarker in clinical research (Figure 4), which allows to identify patients with a tendency of developing a cardiovascular pathology. In conclusion, both platelet stiffness and shape changes are biological parameters that could be exploited as biomarkers in clinical research. Future studies should include patients with a broad range of diseases since platelet activation responses and pharmacological responses are likely to vary according to disease setting.

## Acknowledgments

This work was funded by the Deutsche Forschungsgemeinschaft (DFG, German Research Foundation) – Projektnummer 335549539/GRK2381 and Projektnummer 374031971 - TRR 240. The Facility for Imaging by Light Microscopy (FILM) at Imperial College London is partly supported by the Wellcome Trust (grant 104931/Z/14/Z) and BBSRC (grant BB/L015129/19). A.B. is supported by the Add-on Fellowship of the Joachim Herz Foundation.

## Informed Consent Statement

Informed consent was obtained from all the subjects involved in the study in Tübingen and London.

## Author Contribution

Conceptualization: J.R., A.B., J.S., D.P.Q., M.E., F.S., R.F., S.F., and T.E.S; Investigation: J.R., A.B., J.S., D.P.Q., and A.K.; Resources: A.F., M.E., M.B., S.F., R.F., and T.E.S.; Writing – original draft preparation: J.R; Writing - review and editing: J.R., A.B., J.S., D.P.Q., A.K., M.B., M.F., A.F., M.E., F.S., R.F., S.F., T.E.S.; Visualization: J.R., A.B., J.S., D.P.Q., A.K., M.B., M.F., A.F., M.E., F.S., R.F., S.F., T.E.S.; Supervision: M.E., F.S., R.F., S.F., T.E.S.; Project administration: J.R., T.E.S.; Funding acquisition: T.E.S. All authors have read and agreed to the published version of the manuscript.

